# evoCancerGPT: Generating Zero-Shot Single-Cell and Single-Sample Cancer Progression Through Transfer Learning

**DOI:** 10.64898/2026.02.12.705621

**Authors:** Xi Wang, Runzi Tan, Simona Cristea

## Abstract

Cancer evolution is driven by complex changes in gene expression as cells transition and change states during tumorigenesis. Single-cell RNA sequencing has provided snapshot insights into how the transcriptomics of tumors evolve, but whether the existing knowledge can be used to reliably learn and generate the patterns behind the evolution of cancers remains unknown. Here, we introduce evoCancerGPT, a generative pre-trained transformer decoder-only single-cell foundation model designed to forecast future gene expression profiles in cancer evolution by leveraging previous cell states at the level of single patients. This model integrates the continuous gene expression data of each cell to create a comprehensive representation of a cell token. Training sentences are constructed for each cancer type, each patient and each cell type separately, ordered via inferred pseudotime algorithms, using 2.76 million cell tokens, each with 12,639 genes, spanning 7 cancer types. By learning from long-range dependencies between cells arranged in pseudotime from a large corpus of data, evoCancerGPT captures key transitions in cancer evolution, achieving high concordance to ground truth trajectories and outperforming linear and scGPT baselines in held-out test samples in low-context scenarios. Our work suggests evoCancerGPT’s potential utility in characterizing tumor progression at a single-cell and single-patient level and ultimately contributing to more personalized cancer care.

## 1 Introduction

Cancer progression involves a series of intricate genetic and epigenetic alterations that drive the transformation of normal cells into malignant ones. These changes are marked by complex shifts in gene expression profiles as cells transition through various stages of tumor development, from normal tissue to precursor lesions, primary tumors, and eventual metastasis [Hanahan and Weinberg, 2011]. Understanding these dynamic processes at the single-cell level is crucial, as it unveils the heterogeneity within tumors and the evolutionary paths that cancer cells undertake.

Single-cell RNA sequencing (scRNAseq) has emerged as a powerful tool to dissect the transcriptional landscape of tumors at unprecedented resolution [Tang et al., 2009, Navin et al., 2011]. By profiling individual cells, scRNAseq enables the identification of distinct cellular subpopulations, including rare cell types that may drive disease progression or therapy resistance [Patel et al., 2014]. Importantly, scRNAseq data can be leveraged to reconstruct developmental trajectories using computational methods such as pseudotime or trajectory analysis, providing insights into the temporal order of cellular states during tumor progression [Trapnell et al., 2014, Haghverdi et al., 2016, Setty et al., 2019]. Traditional pseudotime approaches, while valuable, are inherently retrospective: they infer past cellular transitions based on snapshot data from heterogeneous cell populations [Saelens et al., 2019]. These methods often rely on assumptions regarding the underlying tumor data or its expected properties that may oversimplify the true biological complexity of cancer progression. Moreover, existing approaches do not inherently possess predictive capabilities to forecast future cellular states or simulate tumor evolution under different or novel conditions.

Recent advancements in artificial intelligence, particularly in natural language processing, have introduced transformer-based models capable of capturing long-range dependencies within sequential data [Vaswani et al., 2023]. These models have shown remarkable success in generating coherent and contextually relevant text by learning from large corpora. Translating this success to biological data presents an intriguing opportunity: can we apply similar generative modeling techniques to predict and simulate cancer progression at the single-cell level and unique to each patient?

In this work, we present evoCancerGPT (Figure 1), a transformer-based generative model designed to forecast future gene expression profiles of cancer cells by harnessing scRNAseq data. Leveraging a decoder-only transformer architecture with 17.7 million parameters, evoCancerGPT treats each cell’s gene expression profile as a token within a sequence ordered by pseudotime, while also allowing for bifurcations or multiple branches in the underlying ordering, with each branch represented as a separate sentence. By training on 2.76 million cells from 699 patients in 7 different cancer types from the CellXGene Consortium [Program et al., 2023], the model learns to generate plausible future cellular states in a zero-shot manner, without additional fine-tuning or supervised labels.

**Figure 1.**
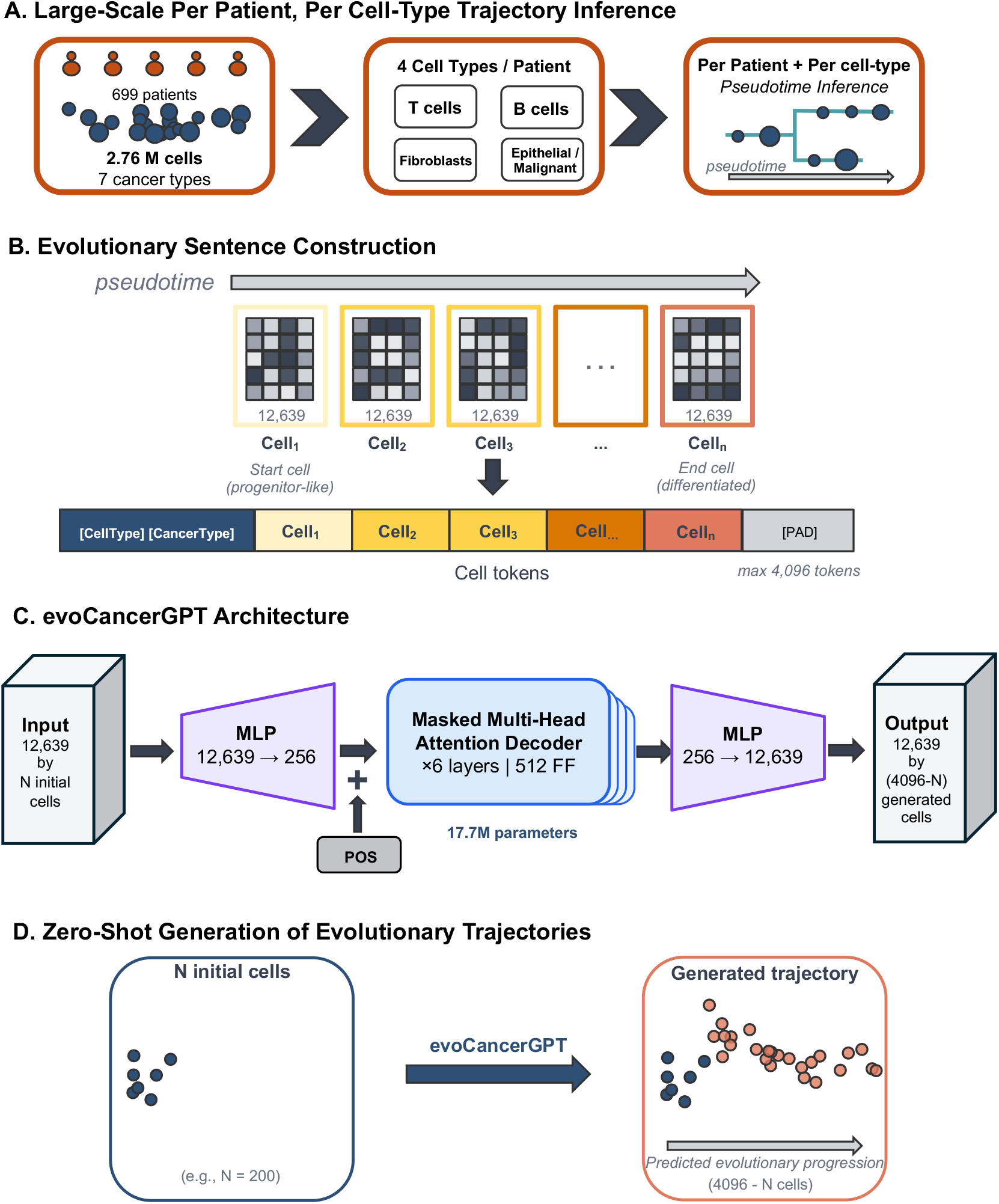
Overview of evoCancerGPT: data preprocessing pipeline (A), detailed sentence constructions (B), architecture (C), and an example inference for next cell prediction given 200 initial cells to generate 3,896 cells (D). Evolutionary sentences are constructed from 4,096 tokens (representing 4,096 or less cells, with the empty space filled with padding up to the 4,096 tokens) in increasing pseudotime order. evoCancerGPT has a decoder-only transformer architecture with an extra MLP layer before and after the transformer blocks to account for the sparse expression matrices and non-linear relationships within scRNAseq data. evoCancerGPT conducts zero-shot generations on the next cells in evolutionary trajectory progressions, given the expression of past progression cells.

Our approach offers several advantages over existing methods. First, evoCancerGPT integrates the continuous gene expression data into the transformer framework, rather than binned values, as done by many of the existing single cell foundation models [Yang et al., 2022, Cui et al., 2024, Szalata et al., 2024]. It therefore captures nuanced transitions between critical evolutionary cellular states. Second, the generative nature of the model allows for the simulation of future trajectories past the point of data collection, providing the option to interrogate cancer evolution into the future. Third, unlike most single-cell foundation models that treat each gene within a cell as a token and each cell as a sentence independent of other cells[Yang et al., 2022, Cui et al., 2024, Szabo et al., 2019, Theodoris et al., 2023], evoCancerGPT treats each cell as a token and all cells from a sample as a sentence, effectively integrating cell-cell dependencies during generations.

We demonstrate that evoCancerGPT not only reconstructs cellular trajectories aligned with observed tumor progression, but it also generates novel gene expression profiles that follow the biology of well-established progression stages. The model successfully captures the progression of key immune and cancer biomarkers, generating cells with expression dynamics that reflect the expected differentiation across cell states. These capabilities suggest that evoCancerGPT may allow for the exploration of hypothetical tumor evolution scenarios, aiding in the identification of potential therapeutic targets and biomarkers for personalized cancer care.

By bridging the gap between generative AI models and evolutionary tumorigenesis at the single-cell level, evoCancerGPT opens new avenues for computational oncology. It provides a framework for simulating and better understanding cancer progression with unprecedented resolution and offers insights that may be unattainable through traditional computational, theoretical, or mechanistic cancer genomics methodologies.

## 2 Model

### 2.1 evoCancerGPT overview

We introduce evoCancerGPT as the first patient-level foundation model trained on cancer trajectories inferred from sc-RNAseq cancer datasets (Figure 1). evoCancerGPT generates zero-shot progression trajectories as novel cells, following an initial number of cells organized as a starting trajectory to be expanded. In our application, the starting trajectory is obtained via pseudotime algorithms (more specifically, Palantir [Setty et al., 2019]); however, any trajectory-generating algorithm or ordering that is considered suitable for the application at hand can be employed, provided that the ordering remains consistent among the sentences used for training. The same ordering will be learned, internalized, and extended by evoCancerGPT with the generation of each new token. evoCancerGPT’s tokenizer is novel: single-sample (here, one sample = one patient) pseudotime-ordered evolution sentences are constructed with each cell’s expression vector of 12, 639 genes as a single token.

### 2.2 Generation of single-sample evolutionary sentences

Each sample’s preprocessed scRNAseq data is categorized into 4 cell types, and sentences are constructed for each patient and each cell type separately: *T cells, B cells, epithelial malignant cells*, and *fibroblasts*. Each sentence represents a singular branch trajectory through pseudotime for a particular cell type. The superset of all the sentences for that cell type represents the entire observed space of possible evolutionary trajectories in gene expression values used for model training. Pseudotime ordering is inferred by running Palantir [Setty et al., 2019] separately for each sample and each cell type. The inferred branching information (sometimes a branching structure, rather than a single 1D linear progression) is used to construct one sentence per branch, independent of the other branches. The maximum context length of each sentence is set to 4, 096. For those branches with over 4, 096 cells, multiple sentences are constructed independently through chunking without overlaps. For more details about how evoCancerGPT generates the evolutionary sentences, see Appendix A and B.

### 2.3 Data and model training

Following pre-training, evoCancerGPT performs zero-shot generation of full expression values for all genes in each generated cell. Fine-tuning with given datasets is also supported; however, it is not exemplified here. For pre-training, we first pre-process sample-level scRNAseq data from a collection of 2.76 million cells from 699 tumor samples of patients with 7 cancer types from the CELLxGENE collection [Program et al., 2023] (see Appendix A). We initially split samples 95%/5% into training and validation sets per cancer type. However, after quality control, preprocessing, and pseudotime inference, we performed a unified donor-level resplit across all cancer types to ensure globally consistent donor assignments before sample-level and cell-type-level sentence construction (see Appendix A.7).

### 2.4 Model architecture

evoCancerGPT uses a decoder-only transformer architecture with two extra layers: one multilayer perceptron layer (MLP) before the transformer blocks for projecting the entire expression matrices of cells into dense embeddings, and another MLP layer after the transformer blocks that projects the output embeddings of the transformer block back to the expression matrix (Figure 1). This allows the model to be trained with sparse matrices of scRNAs-seq expression and to output the continuous expression values of all genes for each newly generated cell. In total, evoCancerGPT utilizes 6 transformer layers, with around 17.7 million parameters (for more details, see Appendix B). The pre-training loss function is the mean squared error (MSE).

## 3 Results

### 3.1 evoCancerGPT generates reliable single-sample and single-cell cancer trajectories in a zero-shot setting

To evaluate whether evoCancerGPT can accurately learn the underlying principles of cancer progression, we investigated its ability to generate sample-level cells across each cell type and cancer-type pair belonging to patients who were completely unseen during pre-training (no cells from these patients were included in pre-training; each sample belonged to either training or validation). We initialized the model with different numbers of initial cells (*N*) in the held-out samples: 200, 500, and 1, 024, each representing low (4.9%), medium (12.2%), and high (25%) percentages of the entire 4, 096 context length. We visualized the initial cells, ground truth cells, and generated cells on UMAPs of all genes, with examples shown in Figure 2A-B.

**Figure 2.**
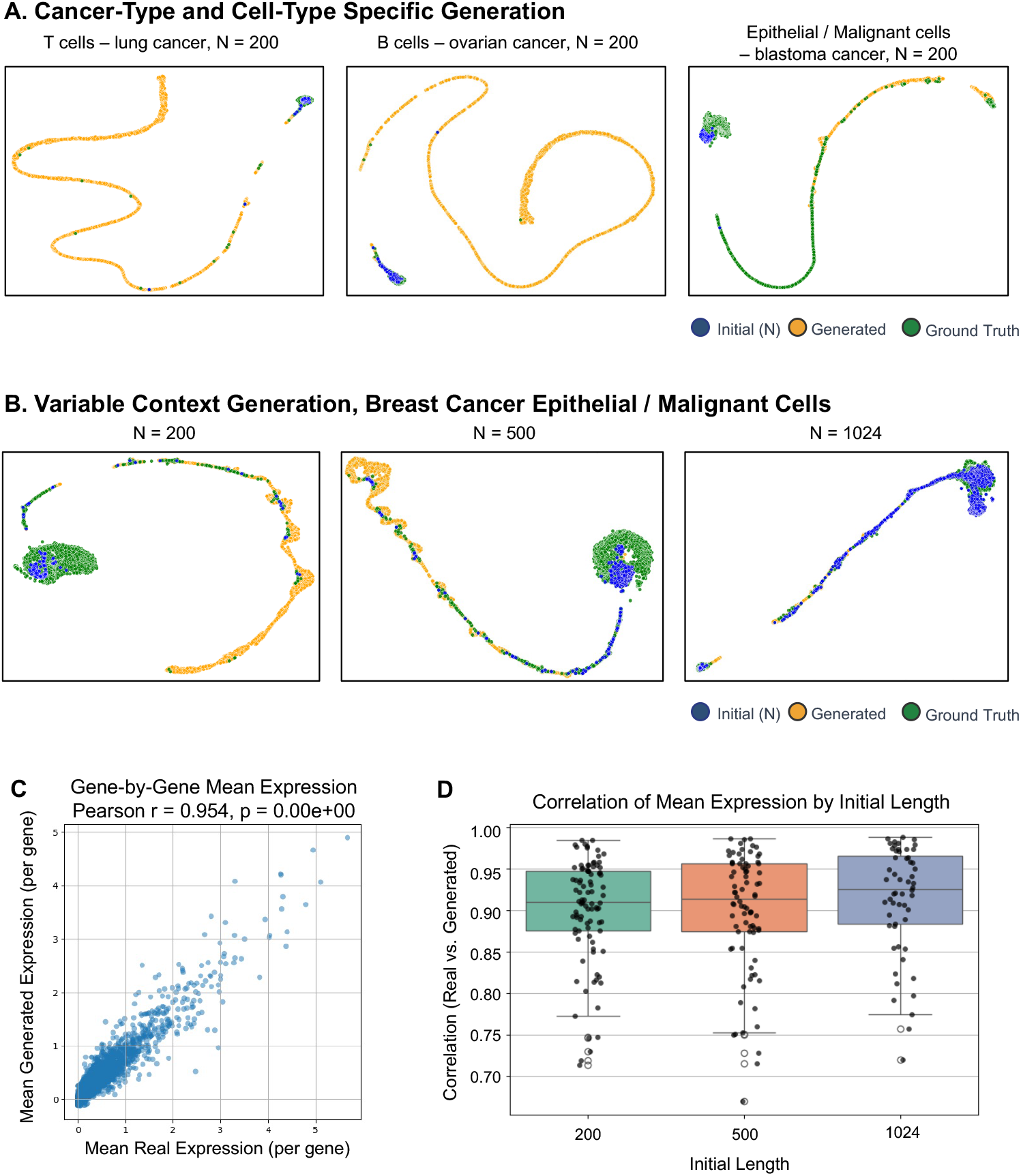
Initial cells, generated cells and ground truth cells across tumor progression, as assessed in validation samples of unseen patients. (A) Sample UMAPs of generated held-out validation sentences across cancer types and cell types with *N* = 200 initial cells. (B) Example variable context generation for breast cancer epithelial/malignant cells across varying numbers of initial cells. (C) Pearson correlation of the mean per-gene expressions of generated vs. ground truth cells given *N* = 200, in one selected sample. (D) Pearson correlation of mean per-gene expressions of generated vs. real cells across all validation samples across varying numbers of initial cells.

We observed that in most of the validation samples across each cell type and cancer-type pairs, the generated cells accurately reconstructed the distributions of the ground truth cells. evoCancerGPT was able to generate cells that fall outside the distributions of the initial cells and match the more differentiated ground truth cells along the entire tumor progression trajectory. If the ground-truth sentences consist of fewer than 4,096 cells, the model will generate more cells than the ground-truth sample, as its architecture requires each inference to fill in the entire 4, 096 cells. These over-generated cells can be seen as potentially more differentiated and advanced cellular states along the tumor progression spectrum than those contained in the sequenced sample.

evoCancerGPT was also able to connect the ground truth cells and the generated cells in different regions of the UMAP, consistently across different numbers of initial cells *N* (Figure 2A-B). The model performed well with just 4.9% of the sample cells when tasked with generating the rest of the 95.1% increasingly differentiated cells across the tumor progression trajectory. Increasing the number of given initial cells constrained the model’s hallucinations and focused the model on generating only along certain regions of the trajectories.

To quantitatively evaluate the similarity between generated and ground truth cells across various numbers of initial cells, we calculated the Pearson correlation coefficient between each of the 12, 639 genes’ mean expressions across the generated and ground truth cells. We found strong correlations between the mean generated expression per gene and the mean real expression per gene across all the validation samples with different numbers of initial cells, demonstrating evoCancerGPT’s ability to accurately reconstruct the overall gene expression patterns of cells along tumor progression trajectories (Figure 2C-D).

### 3.2 evoCancerGPT outperforms baselines in low-context prediction

To benchmark evoCancerGPT against alternative approaches, we compared it with a linear baseline and an scGPT embedding baseline [Cui et al., 2024] using binned MAE (bMAE) and binned Pearson correlation (bPearson), computed on the top 100 pseudotime-associated genes (see Appendix C for details). We also performed a hyperparameter sensitivity analysis evaluating multiple evoCancerGPT (ecGPT) variants with different dropout rates (d) and training durations (ep): ecGPT (0.4 drop 20ep), ecGPT (0.4d, 10ep), and ecGPT (0.1d, 10ep).

At *N* = 200 initial cells, evoCancerGPT variants achieved the lowest bMAE and highest bPearson across held-out validation samples, outperforming both baselines (Figure 3A). The ecGPT (0.4d, 20ep) variant showed the most consistent performance with the least variance. As the number of initial cells increased (Figure 3B), the linear baseline improved, overtaking evoCancerGPT when given *N* = 1, 024 initial cells. This suggests that, according to these two metrics, the transformer architecture is advantageous in low-context settings, while a linear baseline achieves an overall better Pearson correlation and a lower MAE for larger contexts. Similar observations on correlation-like metrics benchmarking transformers against linear models for scRNA-seq data have been documented previously [Ahlmann-Eltze et al., Csendes et al., Wong et al., 2025, Wei et al.], with other reports debating the biological relevance of linear correlations [Miller et al., 2025]. The scGPT embedding baseline showed relatively stable but lower performance across all context lengths, indicating that pretrained single-cell embeddings from a foundation model alone are insufficient to capture the sequential dependencies in cancer progression trajectories.

**Figure 3.**
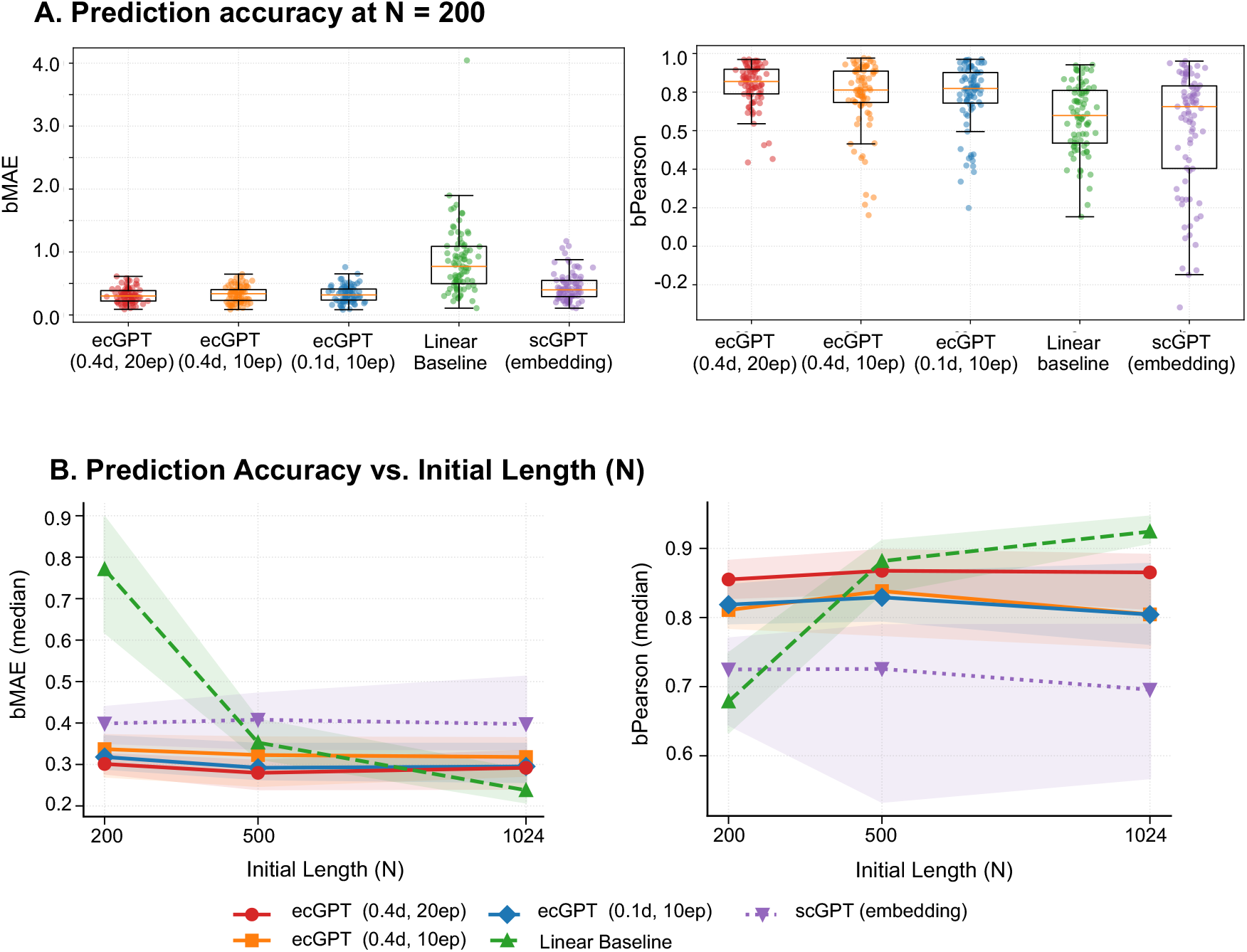
Quantitative evaluations of generated cells vs. ground truth cells. (A) Prediction accuracy at *N* = 200 measured by binned MAE (bMAE, lower is better) and binned Pearson correlation (bPearson, higher is better), comparing evoCancerGPT (ecGPT) variants, a linear base-line, and scGPT embeddings. Metrics are computed on the top 100 pseudotime-associated genes per sentence. (B) Prediction accuracy measured by bMAE and bPearson for varying numbers of initial cells.

### 3.3 evoCancerGPT generates accurate immune and epithelial progressions through key gene programs

The dynamics of cancer progression is marked by known changes in key biomarker genes and gene programs. We further investigated evoCancerGPT’s ability to capture the dynamic expression patterns of tumor progression by assessing whether the generated cell-type and cancer-type specific expression patterns align with literature-documented tumor progression patterns (see Appendix D).

For cancer T cells (T cells sequenced within the cancer samples in our data), we scored literature-documented tumor progression gene sets across the generated cellular trajectories, separately for CD4 and CD8 T cells [Szabo et al., 2019] (Figure 4). Known progression stages for CD4 T cells include Naive/Central Memory, Early Activation, Intermediate (IFN), and Full Activation; while for CD8 T cells, such stages include Naive/Resting, Early Effector, Active Effector, and Full Effector. For the marker genes composing these gene programs, see Appendix D.1.

**Figure 4.**
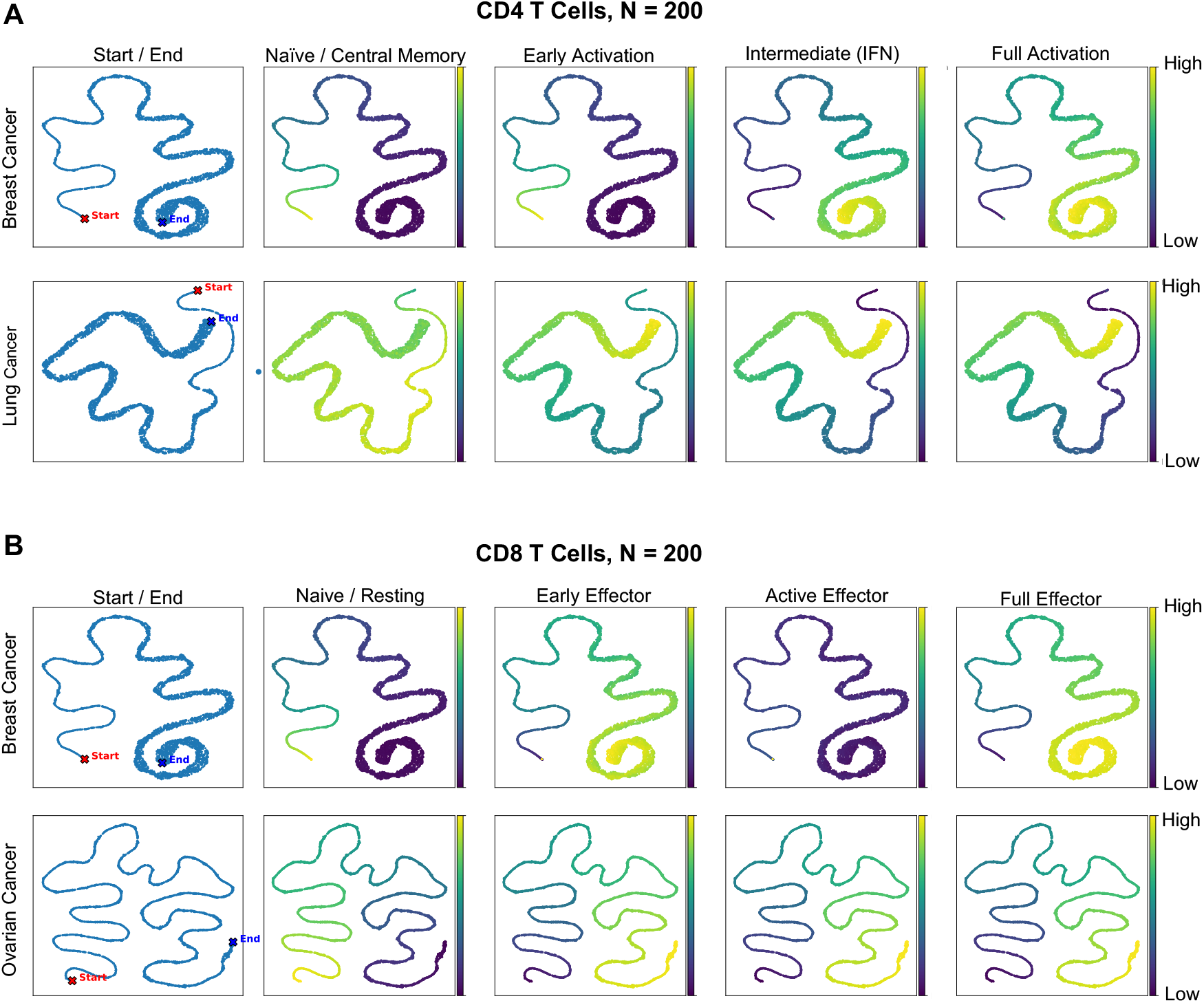
UMAPs of generated CD4 T cells (A) and CD8 T cells (B), colored by their score on literature-documented progression genesets, with initial length *N* = 200, for one held-out unseen patient sample for breast, lung and ovarian cancers. Start and end points of the generated trajectories are marked (see Appendix D.1 for the T cell progression genesets).

evoCancerGPT generated T cells with increasing activation, with the early-generated cells (close to “Start” in Figure 4) showing more progenitor-like gene expression cell states, and the cells generated in later orders showing more differentiated expression and activated cell states (close to “End” in the plots). We exemplified these observations by showing CD4 T cell progression patterns in breast and lung cancer, as well as CD8 T cell progression patterns in breast and ovarian cancer.

We observed similar progression patterns across cell state progression for epithelial malignant cells in breast and lung cancer, using scoring of cancer-specific genesets indicating different stages of tumor progression (Figure 5 and Appendix D.2). The earlier-generated cells expressed earlier epithelial tumor cell state markers, with the later cells in the order of sentence construction expressing intermediate or more advanced markers across cancer types.

**Figure 5.**
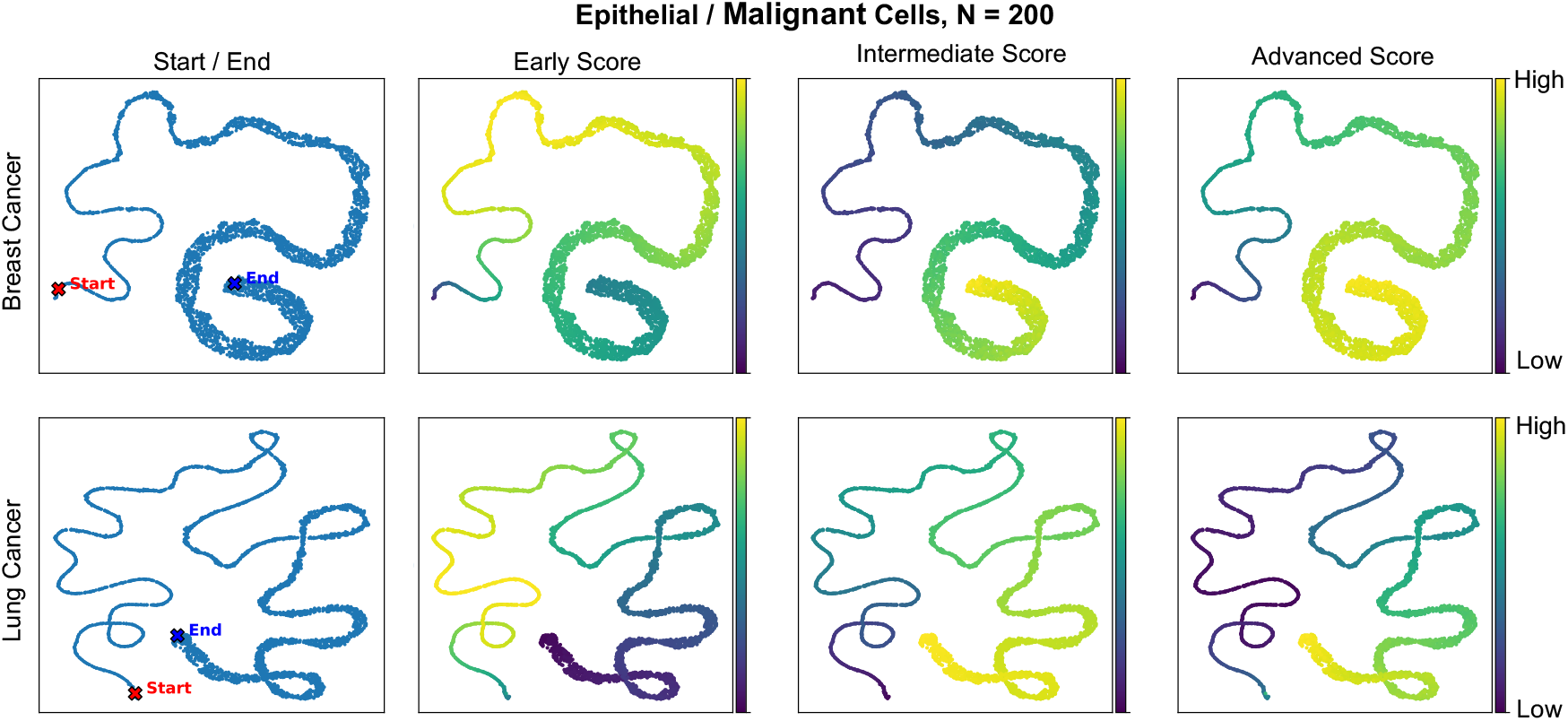
UMAPs of generated Epithelial Malignant cells, colored by their score on literature-documented cancer-specific progression genesets, with initial length *N* = 200, for one held-out unseen patient sample for breast and lung cancers. Start and end points of the generated trajectories are marked (see Appendix D.2 for the tumor malignant progression genesets).

## 4 Conclusion and Discussion

In this study, we introduced evoCancerGPT, a transformer-based generative model designed to simulate evolutionary cancer progression using scRNAs-seq data. By treating each cell’s gene expression profile as a token within sequences ordered by pseudotime, which we term evolutionary sentences, evoCancerGPT learns to generate biologically meaningful future progression cellular states in a zero-shot manner. This approach leverages the capabilities of decoder-only transformer architectures to capture long-range dependencies and complex patterns within high-dimensional single cell omics data.

Our results show that evoCancerGPT can generate cells that follow an increasingly differentiated cancer trajectory in pseudotime, closely mirroring the progression observed in patient scRNAs-seq cancer datasets. The model effectively captures the progression of key cancer biomarker genesets across different cell types and cancer types, producing smooth expression trajectories that align with the general trends in the ground truth data. Notably, evoCancerGPT is able to denoise the inherent variability in single-cell data with respect to cancer progression, suggesting its potential utility in highlighting underlying biological signals of tumor development.

The ability of evoCancerGPT to generate future gene expression profiles based on previous cell states offers a novel perspective in modeling tumor evolution. Traditional pseudotime methods primarily focus on reconstructing past trajectories from snapshot data, lacking predictive capabilities for forecasting future cellular states. In contrast, evoCancerGPT provides a framework for simulating how cancers evolve over time, which is valuable for understanding tumor progression mechanisms and exploring hypothetical scenarios in cancer development.

However, our work is a proof-of-concept study. evoCancerGPT’s outputs reflect extensions of inferred trajectory orderings rather than predictions of true biological time. The generated cells represent plausible continuations along the manifold captured by pseudotime, not verified forecasts of future tumor states in vivo. Pseudotime ordering might not always capture the true complexities of tumor evolutionary time, and it is unclear whether pseudotime truly represents a good proxy for the timing of cancer progression. Therefore, the quality of evoCancerGPT’s outputs is inherently bounded by the quality of the underlying ordering used for training. Nevertheless, our framework is agnostic to the underlying input ordering used for training, and researchers may substitute pseudotime inference with any suitable trajectory algorithm or experimental method capable of generating a reliable estimate of evolutionary time, such as time-series experimental data. In addition, experimental time-series data, which is usually limited in size, could be used to fine-tune cancerEvoGPT to generate more accurate cancer progression trajectories.

In conclusion, evoCancerGPT represents a step towards leveraging generative AI models in cancer research. Our findings suggest that transformer-based architectures can capture essential aspects of cancer evolution and potentially offer valuable insights into tumor progression. Continued development and validation of such models will contribute to personalized cancer care and novel therapeutics.

## 5 Limitations and Ethical Considerations

### Limitations

evoCancerGPT relies on pseudotime inferred from Palantir as a proxy for true evolutionary time, which may not fully capture the complexities of tumor progression in vivo. The model has not been validated on prospective time-series data or against experimentally measured temporal orderings. When the inferred trajectory contains multiple branches, each branch is treated as an independent sentence with no explicit encoding of shared ancestry.

### Ethical considerations

All scRNAseq data used in this study were obtained from the CELLx-GENE Consortium [Program et al., 2023]. No new human samples were collected, and no patient-identifiable information was obtained or used. No institutional review board (IRB) approval was required. evoCancerGPT is a research tool for the computational exploration of cancer progression and is not intended for clinical decision-making or patient diagnosis.

## 6 Code Availability

All codes used for data preprocessing, pseudotime inference, sentence construction, model training, sentence generation, and metrics calculation used to generate figures in this study are open-sourced and publicly available at https://github.com/cristea-lab/evoCancerGPT_main.git

## A Data

### A.1 Data collection

In this study, we utilized single-cell RNA sequencing (scRNAseq) datasets from CELLxGENE [Program et al., 2023].

### A.2 Data loading and initial processing

We loaded all the cancer datasets using Scanpy (v1.10.2) [Wolf et al., 2018].

### A.3 Cell type mapping

We only retained T cells, B cells, fibroblasts, and epithelial/malignant cells. To harmonize cell type annotations across datasets, we mapped the granular cell types provided by the original studies to these four categories.

### A.4 Data filtering and normalization

For each cell type (of the 4 major cell types), we processed single-cell RNA-seq data using Scanpy (v1.10.2) [Wolf et al., 2018]. We applied the following steps to each donor separately:

1. Cell Quality Control
  - Minimum genes: Cells with fewer than 500 detected genes were excluded.
  - Mitochondrial content: We annotated mitochondrial genes (prefix “MT-”) and removed cells with *>* 10% mitochondrial content to filter out low-quality or dying cells.
2. Normalization
  - Starting with the raw gene expression, we normalized the total counts per cell to 10, 000 and applied the *log1p* transformation to stabilize variance and approximate a normal distribution of expression values.
  - For each cell type separately, samples with fewer than 500 cells after quality control were excluded from training.

### A.5 Pseudotime inference with Palantir

We employed Palantir (v1.3.3) [Setty et al., 2019] to infer pseudotime trajectories for each major cell type of interest and for each sample separately. The pipeline consisted of the following steps:

1. Highly Variable Genes (HVGs). We identified the top 2, 000 HVGs using the *Seurat v3* flavor in Scanpy (except for glioblastoma, for which we used 3, 000 HVG). In cases where the Seurat v3 flavor failed, we fell back to the standard Seurat flavor on log-normalized data.
2. Dimensionality Reduction. We performed Principal Component Analysis (PCA) with 50 components on the HVGs followed by Diffusion Maps with 15 components to capture the manifold structure of the data. A multi-scale diffusion component space was then determined for downstream analysis.
3. MAGIC Imputation. We applied MAGIC [Van Dijk et al., 2018] on the diffusion components to smooth the expression data and reduce dropout noise.
4. Palantir Trajectory Inference. We used the *run palantir* function with 300 waypoints to compute pseudotime, terminal state probabilities, and entropy measures. The pseudotime root cell was identified by marker gene scoring (see below). All cells within each cell type and each sample were then assigned a continuous temporal ordering.

### A.6 Root cell identification

Given that pseudotime estimation in Palantir requires defining a starting (“root”) cell, we defined the root cell with specific marker genes for each cancer type. The exact marker genes used for each cancer type to identify the root are displayed in Table 1. T cell, B cell, and fibroblast markers were shared across all 7 cancer types, while epithelial/malignant markers were cancer-type specific. For each cell type and each sample, we scored all cells on the relevant marker geneset using `sc.tl.score_genes.` We selected candidate root cells above the 99.5th percentile of the marker geneset score, and among these, we chose the cell with the lowest total UMI count as the root cell, to minimize the chance of choosing a doublet.

**Table 1:**
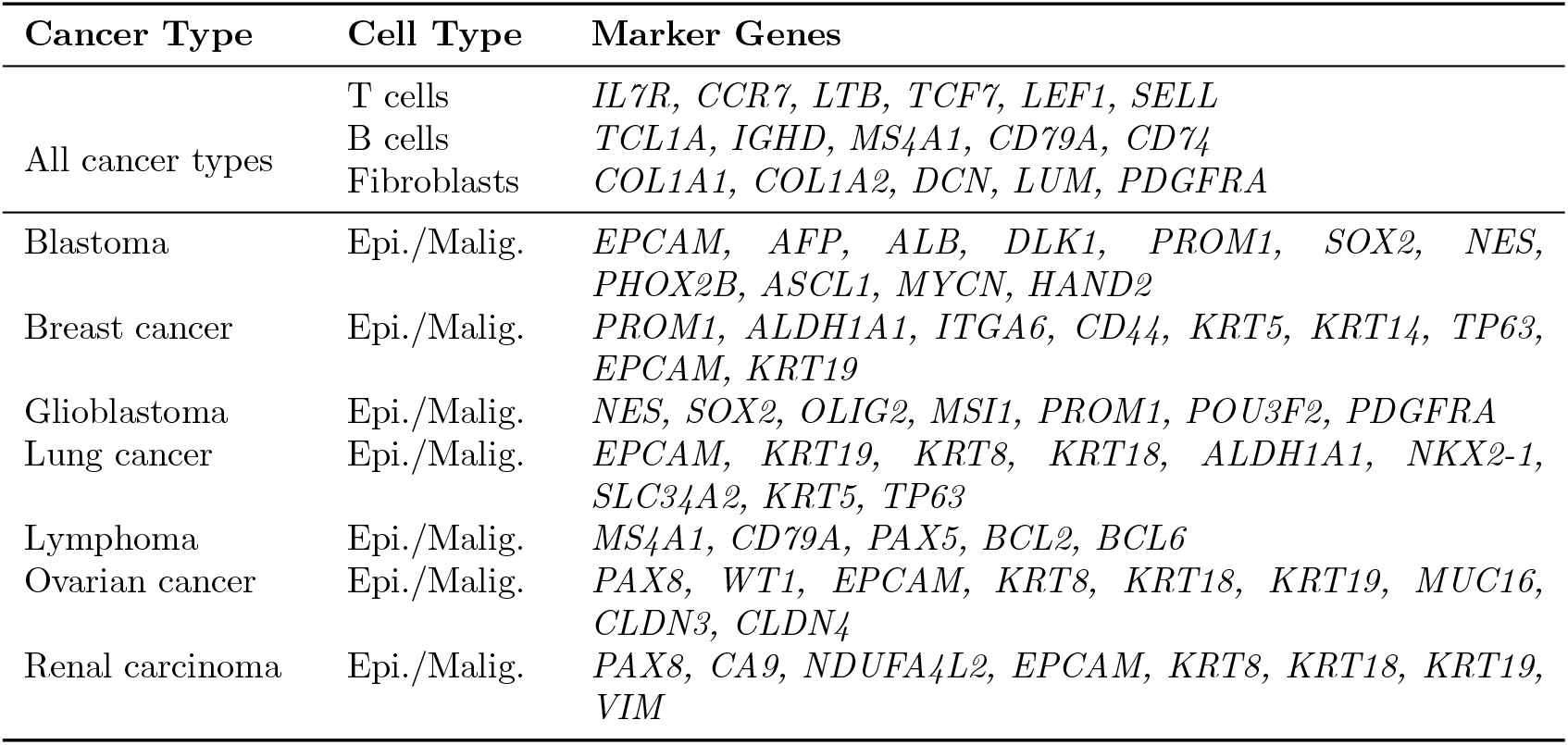
Marker genes used for root cell identification in Palantir, per cancer type and cell type. T cell, B cell, and fibroblast markers are shared across all cancer types. Epithelial/malignant markers are cancer-type specific, reflecting tissue-of-origin biology.

### A.7 Post-pseudotime train/validation resplit

To ensure no data leakage, we performed the initial train/validation split separately for each cancer type before QC. However, after QC, some sample/cell type combinations were removed due to quality or size. Therefore, we performed a donor-level resplit after pseudotime computation and before sentence construction, 95%/5% into training and validation, with the constraint that each cell type has at least three donors in total and at least one donor in the validation set. Cell types with fewer than three donors were assigned entirely to training.

## B Model

### B.1 Data preparation and input representation

#### B.1.1 Cell Tokens

We defined a fixed set of 12,639 genes as the intersection across all samples and used this gene set consistently throughout training and evaluation. Each cell was treated as a single token, represented by its continuous expression vector across all 12,639 genes without binning or discretization.

#### B.1.2 Special Tokens

We prepended seven special tokens at the beginning of each sequence:

1. Cell Type Token (1 token). A learned embedding denoting the cell type (T cells, B cells, epithelial/malignant, or fibroblasts).
2. Cancer Type Token (1 token). A learned embedding identifying the cancer type.
3. Placeholder Tokens (5 tokens). Five learned tokens with fixed indices to be used for custom fine-tuning applications.

### B.2 Model architecture

#### B.2.1 Model components

evoCancerGPT consists of the following components:

1. Gene Embedding MLP. Each cell token (*i*.*e*., cell expression vector of 12,639 genes) was first passed through a two-layer MLP with a GELU activation [Hendrycks and Gimpel, 2023] that projected the vector into an embedding space with 256 dimensions.
2. Cell Type, Cancer Type, and Placeholder Token Embeddings. The cell type and cancer type tokens were each projected to the same 256-dimensional embedding space via separate embedding layers. Five additional placeholder tokens were placed afterward.
3. Positional Embeddings. Learned positional embeddings to represent the position of each token in the sequence were added to the token embeddings.
4. Transformer Decoder Blocks. We used a decoder-only Transformer with 6 layers, 8 attention heads, 256 embedding dimensions, 512 feedforward dimensions, and GELU activations. We trained multiple model variants with dropout rates of 0.1 and 0.4 to evaluate the effect of regularization on generation quality (see Figure 3C,D).
5. Output MLP. After the Transformer layers, we use a two-layer MLP with LayerNorm, GELU, and dropout to project the Transformer hidden embedding back to gene expression space.

#### B.2.2 Training Configuration

We trained evoCancerGPT on a high-performance computing system with one A100 NVIDIA GPU, using the Mean Squared Error (MSE) training objective and the AdamW [Kingma and Ba, 2017] optimizer with 0.05 weight decay (1e-4 learning rate). We used a Batch size of 5, employed ReduceLROnPlateau for Learning Rate scheduling, and trained evoCancerGPT model variants for 10 and 20 epochs. The primary model (ecGPT 0.4d, 20ep) uses a dropout of 0.4 and is trained for 20 epochs.

## C Metrics

### C.1 Quantitative similarity between generated and ground-truth cells

To quantitatively compare the generated cells against ground truth trajectories, we computed two metrics: Binned MAE (bMAE) and Binned Pearson (bPearson). Given *N* initial cells, evoCancerGPT generates cells to fill a total sequence length of 4,096. We compared the generated cells against the corresponding ground truth cells, truncated to the length of the shorter sequence. We excluded sentences where fewer than 256 ground truth cells (after the initial *N* cells) were available.

To emphasize biological signals, we partitioned both generated and ground truth ordered segments into *B* = 20 bins of approximately equal size and averaged gene expression within each bin. To select the subset of genes on which to compute bMAE and bPearson, we computed |Spearman(bin index, truth bin expression)| per gene as a measure of pseudotime association and selected the top *k* = 100 genes with the strongest correlation.

**Binned MAE (bMAE)** is computed as the mean absolute error between the ground-truth and the generated bin-averaged expression across all bins and evaluated genes.

**Binned Pearson (bPearson)** for each bin, we computed the Pearson correlation between the ground-truth and the generated expression across genes. per bin correlations were aggregated using a Fisher z-transform weighted by the degrees of freedom per bin, then back-transformed.

### C.2 Benchmarking baselines

We benchmarked evoCancerGPT against two baselines: a simple linear baseline and a pre-trained foundation model scGPT baseline.

#### Linear baseline

for each evaluation sentence, we fitted a per-gene linear regression on the *N* initial cells (*y* = *β*_0_ + *β*_1_*t*) and extrapolated to generate 4,096 *− N* cells.

#### scGPT embedding baseline

we employed the pretrained scGPT [Cui et al., 2024] whole-human model to encode each cell into a 512-dimensional embedding. We selected 2,000 HVGs per cell-type/cancer-type pair and applied `scg.tasks.embed_data` to obtain cell embeddings. We then trained an evoCancerGPT model with the matching Transformer configuration (6 layers, 256 embedding dim, 8 heads, 512 feedforward dim, 20 epochs, and 0.4 dropout) on the 512-dimensional embeddings. To map the generated embeddings back to gene expression space, we trained a separate MLP (512 → 1024 → 2048 → *n*_genes_, GELU activations, dropout 0.2) on the training data using MSE loss. The MLP was trained with AdamW (learning rate 3 × 10^−4^, batch size 4,096).

## D Cell State Progression Scoring

To assess whether evoCancerGPT generates cells following biologically meaningful progression orders (Figures 4 and 5), we defined marker gene sets for T cell and epithelial/malignant cell state progressions. For each generated validation sentence, we computed a UMAP (15 nearest neighbors) on the full gene expression matrix and scored each cell state as the mean expression across the corresponding marker genes.

### D.1 Immune cell progression markers

#### CD4 T cell states (Figure 4A)

- Naive/Central Memory: *CCR7, SELL, TCF7, LEF1, IL7R*
- Early Activation: *CD69, IL2, TNFRSF4*
- Intermediate (IFN): *IFIT3, IFIT2, STAT1, MX1, IRF7, ISG15*
- Full Activation: *IL32, CCL4, CCL5, IFNG, GZMB, PRF1*

#### CD8 T cell states (Figure 4B)

- Naive/Resting: *CCR7, SELL, TCF7, IL7R*
- Early Effector: *CCL5, GZMK, CXCR6*
- Active Effector: *IFNG, CCL3, CCL4*
- Full Effector: *GZMB, PRF1, NKG7, GNLY, FGFBP2*

### D.2 Epithelial-Malignant progression markers

#### Breast cancer (Figure 5, top)

- Early: *ESR1, PGR, EPCAM, KRT18, CDH1*
- Intermediate (EMT): *SNAI1, SNAI2, ZEB1, TWIST1*
- Advanced: *VIM, FN1, CDH2, MMP2, MMP9*

#### Lung cancer (Figure 5, bottom)

- Early: *NKX2-1, EPCAM, KRT18, CDH1*
- Intermediate (EMT): *SNAI1, SNAI2, ZEB1, TWIST1*
- Advanced: *VIM, FN1, CDH2, MMP2, MMP9*

